# How Mentoring Networks Shape Early-Career Grant Success: Evidence from NIH K-awardees

**DOI:** 10.64898/2026.08.11.743320

**Authors:** Felicia J. Setiono, Eamen Ho, W. Marcus Lambert

**Affiliations:** Department of Epidemiology and Biostatistics, SUNY Downstate Health Sciences University

## Abstract

Effective mentorship is essential for strengthening the STEMM (Science, Technology, Engineering, Mathematics, and Medicine) workforce, yet empirical evidence on how mentorship networks are structured and linked to career success remains limited. Here, we analyze mentorship networks among recipients of NIH career development (K) awards to characterize network size, mentor roles, and their associations with mentee-reported outcomes, including potential variation by sociodemographic characteristics. We found that K-awardees rely on mentors beyond their primary advisor, who play varying roles beyond being a Research mentor. Different mentor roles led to different types of mentoring outcomes; while Research mentors were associated with research-related outcomes such as Publications and Grants, career- and psychosocial-related mentoring outcomes were more likely to come from other types of mentors, such as Coaches, Connectors, and Sponsors. Larger networks, as well as having Peer and Identity mentors are additively beneficial for researchers who identify as underrepresented in science more than their counterparts. This study provides large-scale evidence on how mentorship network configurations relate to early-career grant success.

## Introduction

Mentorship is widely recognized as a cornerstone for advancing the quality and retention of the Science, Technology, Engineering, Mathematics, and Medicine (STEMM) workforce. Numerous studies have shown that effective mentorship is linked to increased productivity, career choice and satisfaction, as well as reduced burnout for individuals working in STEMM settings (1–5). Accordingly, policies and programs that aim to strengthen mentorship have become more commonplace, particularly for early-career researchers (4,6,7). Even certain funding agencies, such as the National Science Foundation (NSF) and the National Institute of Health (NIH), have required formal mentor training for training grants, and structured mentoring plans as components of career development awards applications (8–11).

Despite the formalization of mentorship within federal funding mechanisms, less attention has been given to how mentorship is structured for advanced trainees, such as medical residents and postdoctoral scholars (postdocs), during their transition to career and research independence. This is a critical gap that needs to be addressed as these advanced trainees make up a significant portion of the STEMM workforce (12,13) and are an essential component for retaining scientific talent (14–16). Multiple mentors in the form of a mentorship team or network can be particularly helpful to postdocs, as they tend to have ambiguous mentoring structures during training (14). Unlike graduate students who commonly have a committee of mentors as part of their training, postdocs typically work with one primary mentor (15,16). The burden of choosing the ‘right’ mentor as a postdoc then becomes enormous, since postdocs’ mentors may influence one’s success much more than graduate students’ mentors (17). Postdocs have also reported stress from enormous working hours expectations and limited job prospects (18–21), which are exacerbated by recent funding cuts (22–24). Overall, there is a rising concern for attrition of scientific talent from the United States (U.S.) and effective mentorship networks for postdocs can help buffer the challenges they are facing.

One barrier to developing and formalizing mentoring networks for postdocs is that there remains varying opinions on what kind of mentors and mentoring structure(s) would most effectively support postdocs (17,25). Generally, there is a growing support for cultivating a mentor network, where scientific trainees have multiple mentors who can serve different roles and fulfill trainees’ various needs (e.g., research, career, and psychosocial) (5,26–30). Mentors can provide guidance to develop certain specific skills, serve as an advocate, or focus on increasing one’s social and professional network. These mentors are often referred to as coaches, sponsors, connectors (31,32). The impact of other types of mentors, such as peer mentors, and those who share/affirm identities have also been documented (5,27,33–35). Mentorship relationships can be informal or formal, happening within one’s organization or across external entities, such as professional organizations (36). There is also a push in increasing mentee’s agency in the relationship (‘Mentoring Up’), promoting a more bidirectional and collaborative approach to mentorship relationships between trainees and mentors (7,37). Though it is important to note, that very rarely will the above studies focus on the needs and experiences of postdocs exclusively.

Additionally, trainees of different identities and backgrounds have extremely varying experiences. Racial and ethnic minorities, and others who identify as an underrepresented group in STEMM (UR) face unique challenges during their scientific training (15,38,39) and may require different mentorship structures than their counterparts (40). Some have found, for example, that peer mentors and role models who share, or at least, understand these struggles may be particularly important for UR individuals (28,39,41). Others, stress the importance of mentors who can reinforce science identity for UR individuals (38,42). Continued study in understanding factors that can promote UR postdoctoral transition to faculty is imperative, given the persistent gap in the presence and retaining of UR individuals at the faculty level in academia and medicine, despite increases in the number of UR trainees (43).

Effective mentorship strengthens STEMM trainees and the broader research enterprise by cultivating a well-prepared, connected, and diverse pool of future scientists (1–5). Yet, despite increasing institutional emphasis on structured mentoring, there remains limited empirical evidence characterizing how mentorship networks are configured among advanced trainees and whether these configurations are associated with measurable indicators of career success. Hence, in this study, we examine mentorship networks among researchers who successfully received NIH career development (K)-awards to characterize network size, mentor roles, and associated outcomes. Specifically, our research aims are as follows: 1) identify common structural characteristics of mentoring networks among NIH K-awards recipients, 2) elucidate how distinct mentor roles relate to specific mentee-reported outcomes, and 3) explore how the benefits of multiple mentors may vary across individuals of differing race/ethnicity. By directly capturing both formal and informal mentors beyond the primary advisor across various scientific fields, this study provides one of the first large-scale empirical examinations of how mentorship network configurations relate to early-career grant success.

## Methods

### Experimental Design

The overall objective of this study is to investigate common characteristics of mentorship network among NIH K-Awardees and their association with mentoring outcomes.

#### Participants and recruitment

Individuals who applied for an NIH K-Award (e.g., K01, K08, K22, etc.) between July 1, 2018 – June 30, 2023 were eligible to participate in the study. Participants were recruited through multiple channels; 1) NIH RePORTER was used to collect and tabulate the academic institutions K-Awardees were most associated with. Research deans and Office of Sponsored Programs from these institutions were asked to forward a general email about the study to their postdocs. 2) a list of NIH-supported PIs hosted on the NIH Freedom Information Act Office’s site (44) received individualized emails about the study from the research team. At the time of recruitment, emails of PIs supported between 2018-2022 were available. 3) a paid announcement to postdocs was sent through the National Postdoc Association. 4) Finally, participants of the MOSAIC program (started in 2021) were contacted individually. Names of the program’s awardees were obtained from the NIH RePORTER using PAR-19-343, PAR-21-271, -272, -273 as search terms. Regardless of the recruitment channel, potential participants received a recruitment flyer and a study blurb with a link to the online survey. The survey was open from March – November 2024.

#### Ethical Approval

The study’s consent form was provided to participants online prior to the start of the survey. Participants who consented online were considered enrolled in the study. Those who completed the survey were eligible to enter a raffle for a chance to win one of twenty $50 e-gift cards. The study received an exempt determination from SUNY Downstate IRB office #1981725.

#### Survey

The anonymous survey was hosted online using Qualtrics and took around 30 minutes to complete. The first section of the survey focuses on information about the K-award applications (e.g., type of K-award, award score, etc.) The second section of the survey asks questions regarding the applicants (e.g., demographic information, career outcomes, etc.). The third section makes up the bulk of the survey and asks applicants to list out all the mentors they had during the time of application. The applicant was also asked to indicate any connection between each of the mentors they provided.

### Measures

#### K-award Scores

K-awards are career development awards intended for investigators at earlier phases of their research career to enhance skills development and provide time for research and training activities in the biomedical, behavioral, or clinical sciences. There are multiple “K” Award types designed for different scientific/educational backgrounds and career stages, offered by different NIH institutions, each having different K-programs and requirements. The final impact scores run from 10 – 90, where 10 is the best.

#### Mentoring Roles

For each mentor, the applicant was asked to indicate the mentors’ roles and what the applicant received from the mentoring relationship (Definitions available in Table 1.). In terms of roles, the applicant was allowed to pick from a list of “Research mentor”, “Coach”, “Sponsor”, “Connector”, “Peer mentor”, and “Identity mentor”. Applicants first had to indicate each mentor’s primary role, followed by an option to add other roles that the mentor may play.

**Table 1.**
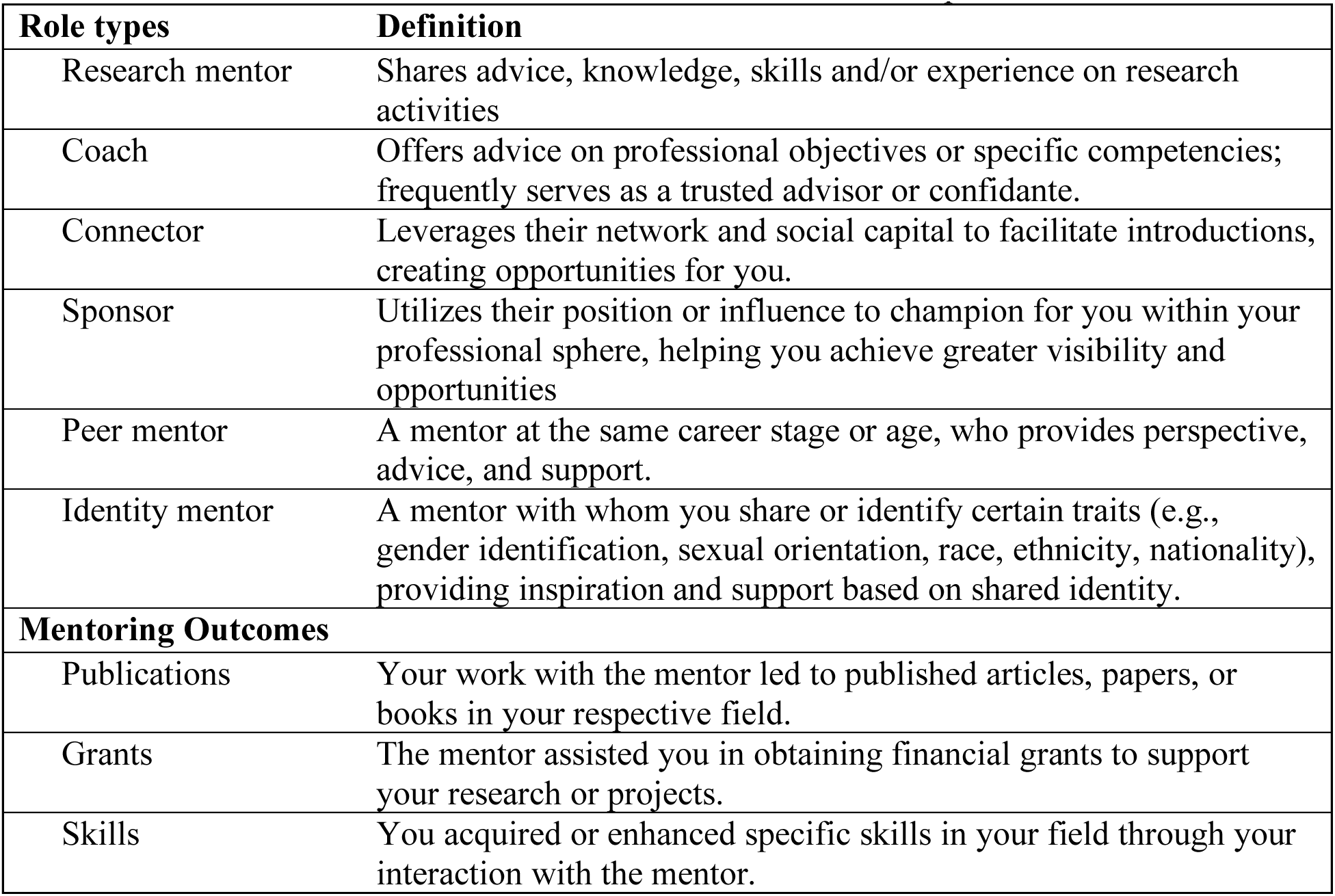
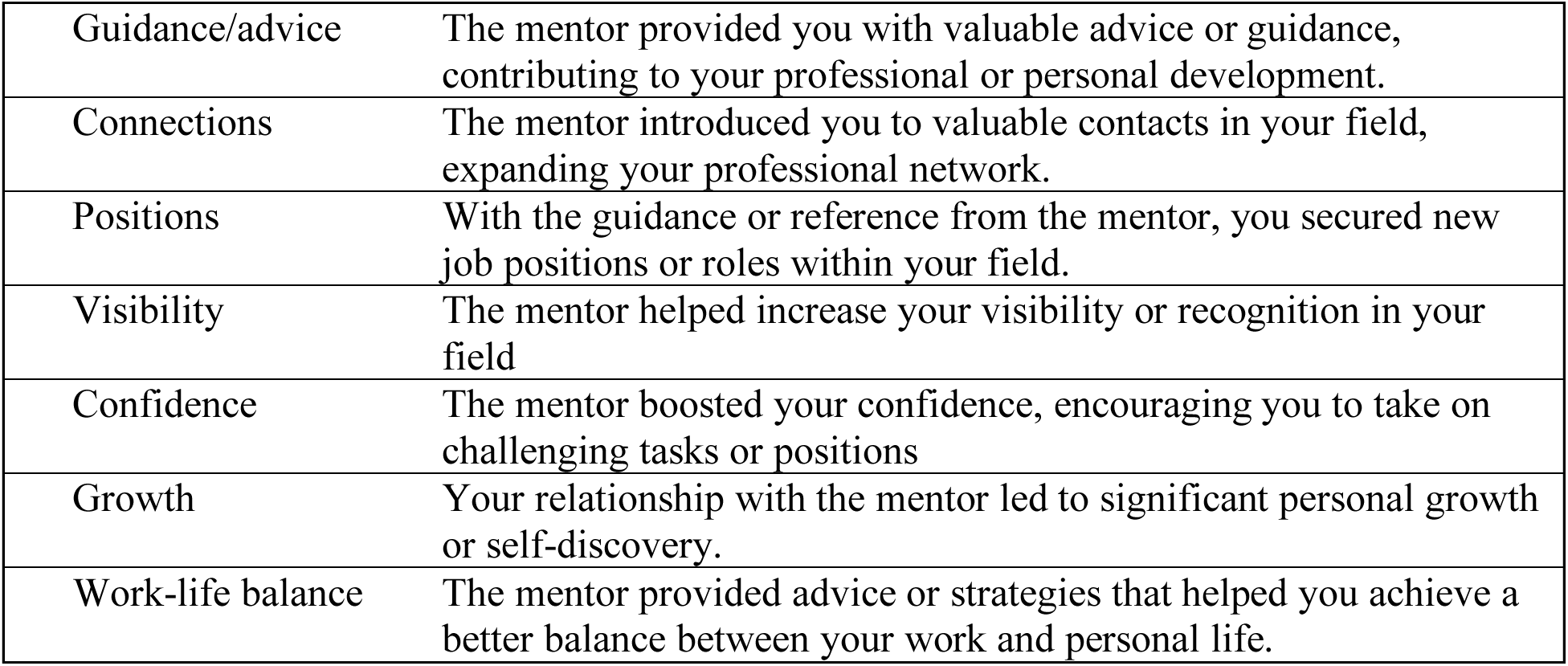
Definitions of all mentor role types and mentoring outcomes included in the survey for K-awardees. Participants were asked to list out their mentors and indicate for each mentor, the mentor’s roles and the outcomes that resulted from the mentorship.

#### Mentoring Outcomes

Applicants were then asked to indicate what they received from the mentoring relationship, henceforth called *mentoring outcomes*. The applicant was allowed to select from the following: “Publications”, “Grants”, “Skills”, “Guidance”, “Connections”, “Positions”, “Visibility”, “Confidence”, “Growth”, “Work-life balance”.

#### Network Size and Density

For each K-awardee, the network’s size and density were calculated. Network size is defined as the number of mentors K-awardees have, while density is calculated as the number of existing connections divided by the total possible connections between all the individuals in the network. The primary role mentors play, and the mentoring outcomes K-awardees receive from their mentors were noted. Within each K-awardee, the number of unique mentor roles was calculated by summing the number of distinct roles their mentors play. Similarly, the number of unique mentoring outcomes was calculated by summing the number of distinct outcomes each K-awardee received from their network.

#### Participant Demographics

Applicants were asked to indicate their current employment sector, field of study, gender, race, and ethnicity, and if they are first-generation low-income background. Due to low numbers, those who identify as Hispanic, Black/African American, Native Hawaiian/Pacific Islander, and American Indian/Alaskan Native were grouped as one category: racial/ethnic underrepresented minority. In this paper’s analysis, this group is labeled as ‘UR’ for underrepresented group.

### Statistical Analysis

A total of 994 individuals completed the online survey. For the purposes of this study, only those who received the K-award (K-awardees) were included in the final analytical sample (N=892). Demographic characteristics of the K-awardees and other descriptive statistics on their applications were calculated. Numerical variables were described using means (SD) or median (range), depending on their distribution. Categorical variables were described using proportions. All analysis was conducted in Python 3.12.4 using NumPy (v 1.26.4), pandas (v 2.2.2), Plotly (v 5.22.0) for visualization, and statsmodels (v 0.14.2) for statistical analysis.

#### Mentoring roles and outcomes

The association between network size and number of distinct mentor roles, as well as number of distinct mentoring outcomes were calculated using multi linear regression models (example model is shown in Eq.1). The models were further adjusted for K-awardees’ demographic characteristics including UR status, gender, and first-generation status.

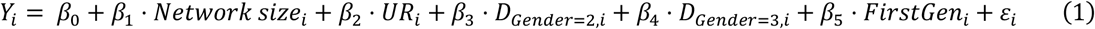

Relationships between types of mentor roles and mentoring outcomes were explored using several multi logistic regression models. In these models, analysis was done at the level of the mentors rather than the K-awardees. Each mentoring outcome was operationalized as a binary variable (whether the mentor resulted in the mentoring outcome) with mentor roles as an independent categorical variable. Example model is shown in Eq.2. Differences in odds ratios of each mentoring outcome being attributed to each mentor role were calculated across all possible roles using pairwise comparisons. Statistical significance was set at a= 0.05 after Bonferroni correction to account for multiple testing done.

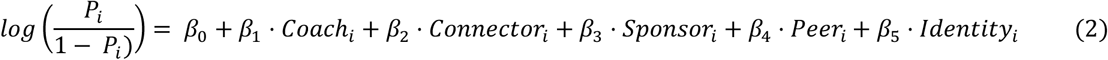

where

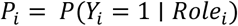

#### K-Award scores and network characteristics

The relationship between K-award scores and network characteristics i.e., network size and density, was examined using a mixed linear effect model with NIH and academic institutions as group level (second-level effects). Moderation of the relationship by race and ethnic categories was investigated.

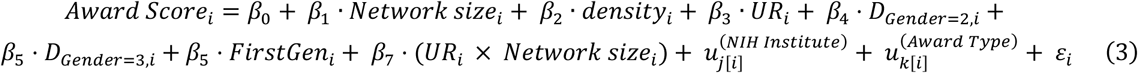

where

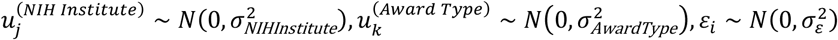

and *j*[*i*]and *k*[*i*]indicate which NIH institute and K-award type observation *i* belongs to, respectively.

Finally, the relationship between K-award scores and mentor roles, as well as mentoring outcomes was examined to identify whether having certain mentor roles are associated with better award scores, with a particular focus on identifying whether these relationships are moderated by UR status. These were also calculated using several mixed linear effect models as above, one for each mentor role and for each mentoring outcome (see Eq. 4. And Eq. 5.) In the model for mentor roles, each role was treated as a binary (i.e., whether the K-awardee has a mentor with that primary role or not). Whereas each mentoring outcome was treated as a numerical variable (i.e., the total number of each mentoring outcome the K-awardees received from their mentors).

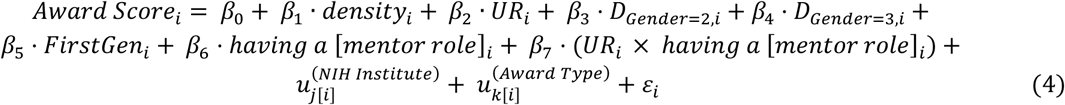

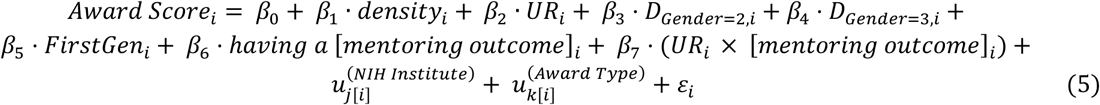

where

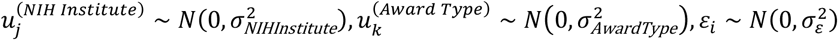

and *j*[*i*]and *k*[*i*]indicate which NIH institute and K-award type observation *i* belongs to, respectively.

## Results

### Characteristics of K-awardees and submitted awards

The sociodemographic characteristics of the final analytical sample (*N*=892) is reported in Table 2. Most of the K-awardees are currently academic faculty members (93.8%) from various fields including Medicine, Behavioral and Social Sciences, and Neuroscience. There were fewer male respondents than female (62.9%) and the majority identified as White (69.1%), followed by Asian (16.5%). The underrepresented group (UR) composed 15.8% of the sample.

**Table 2.** Sociodemographic characteristics of survey respondents who applied and were awarded for an NIH K-award between 2018-2023 (N=892).

| Characteristics | N (%) |
| --- | --- |
| Current employment sector |  |
| Postdoctoral Researcher | 33 (3.7) |
| Academic Faculty (Research-Focused) | 817 (91.6) |
| Academic Faculty (Teaching-Focused) | 20 (2.2) |
| Other Research-Intensive | 13 (1.5) |
| Non-Research, Science-Related | 8 (0.9) |
| Non-Science Related | 1 (0.1) |
| Field of Study ($N=891$) | |
| Behavioral and Social Science | 96 (10.8) |
| Biochemistry | 25 (2.8) |
| Cell and Developmental Biology | 41 (4.6) |
| Engineering | 20 (2.2) |
| Genetics | 18 (2.0) |
| Immunology | 14 (1.6) |
| Medicine | 267 (29.9) |
| Microbiology | 29 (3.3) |
| Molecular Biology | 34 (3.8) |
| Neuroscience | 84 (9.4) |
| Nursing | 21 (2.4) |
| Pharmacology | 28 (3.1) |
| Physiology | 28 (3.1) |
| Public Health | 68 (7.6) |
| Other (<10 responses)* | 118 (13.2) |
| Gender |  |
| Female | 561 (62.9) |
| Male | 322 (36.1) |
| Other/Unknown | 9 (1.0) |
| Race |  |
| Asian | 147 (16.5) |
| Black | 45 (5.0) |
| American Indian, Alaskan Native | 3 (0.1) |
| White | 616 (69.1) |
| Other | 36 (4.1) |
| More than one race | 38 (4.3) |
| Unknown or Not Reported | 7 (0.8) |
| Hispanic | 97 (10.9) |
| First-generation/low-income background | 232 (26.0) |

\*examples of Other fields: Applied Physics, Audiology, Zoology, Translational Medicine, Fisheries, Food Science, etc.

The most common NIH award type submitted was K01 (29.0%), followed by the K23 (28.7%). K-awardees received the award from various NIH Institution including NHLBI, NIDDK, NIMH, NIAID, NCI, and others (Table 3). On average, K-awardees received a score of 22.1 (SD= 7.0) on their application.

**Table 3.**
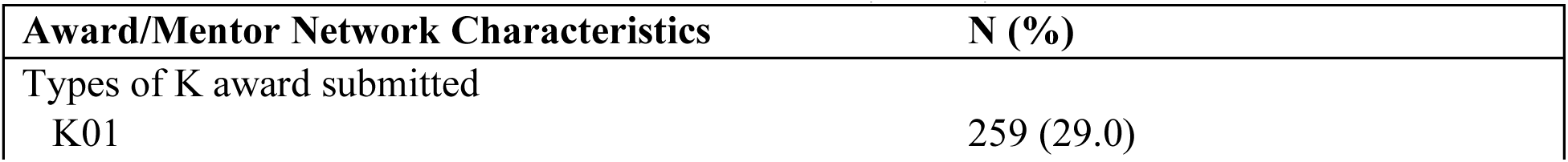
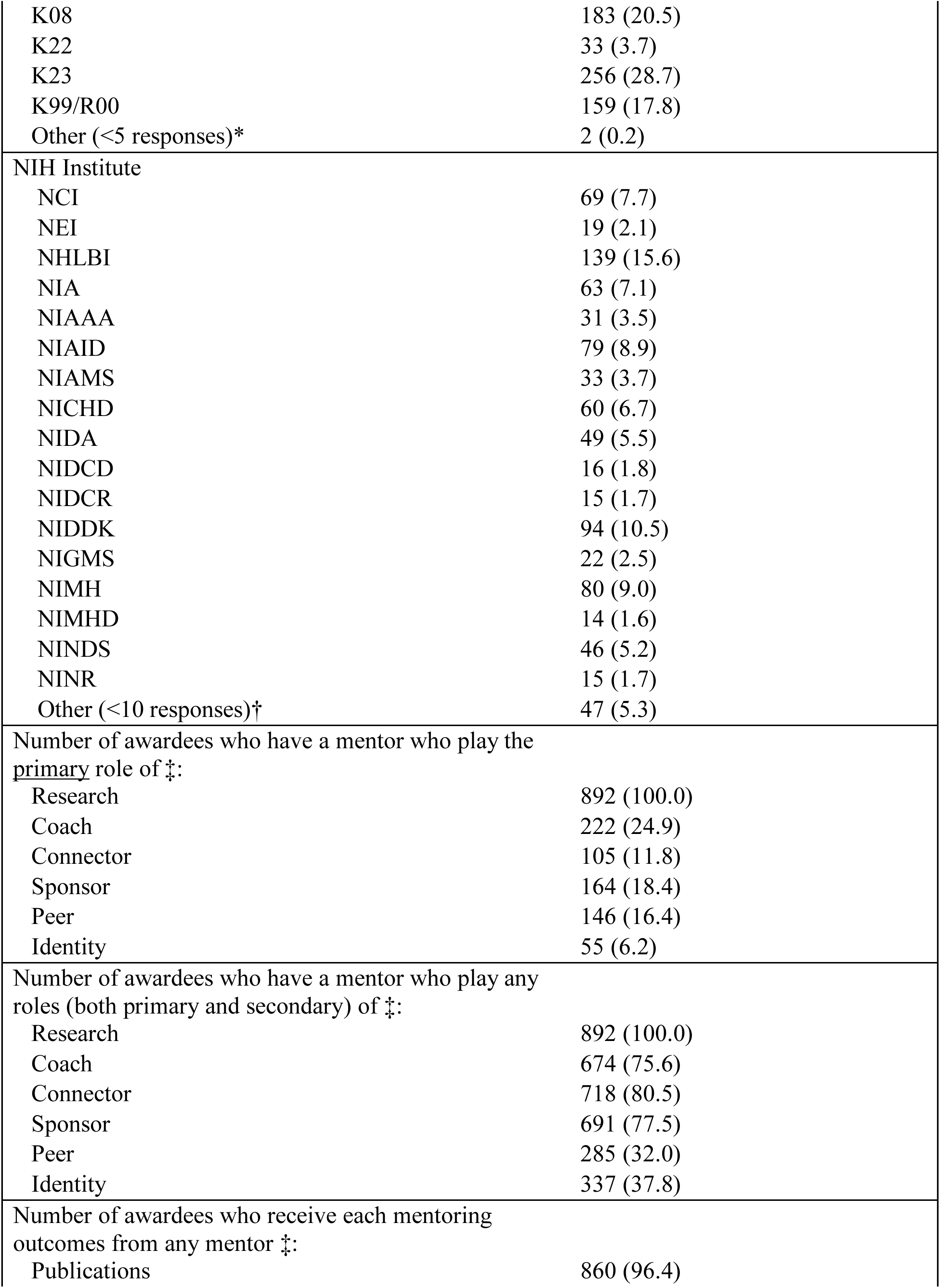
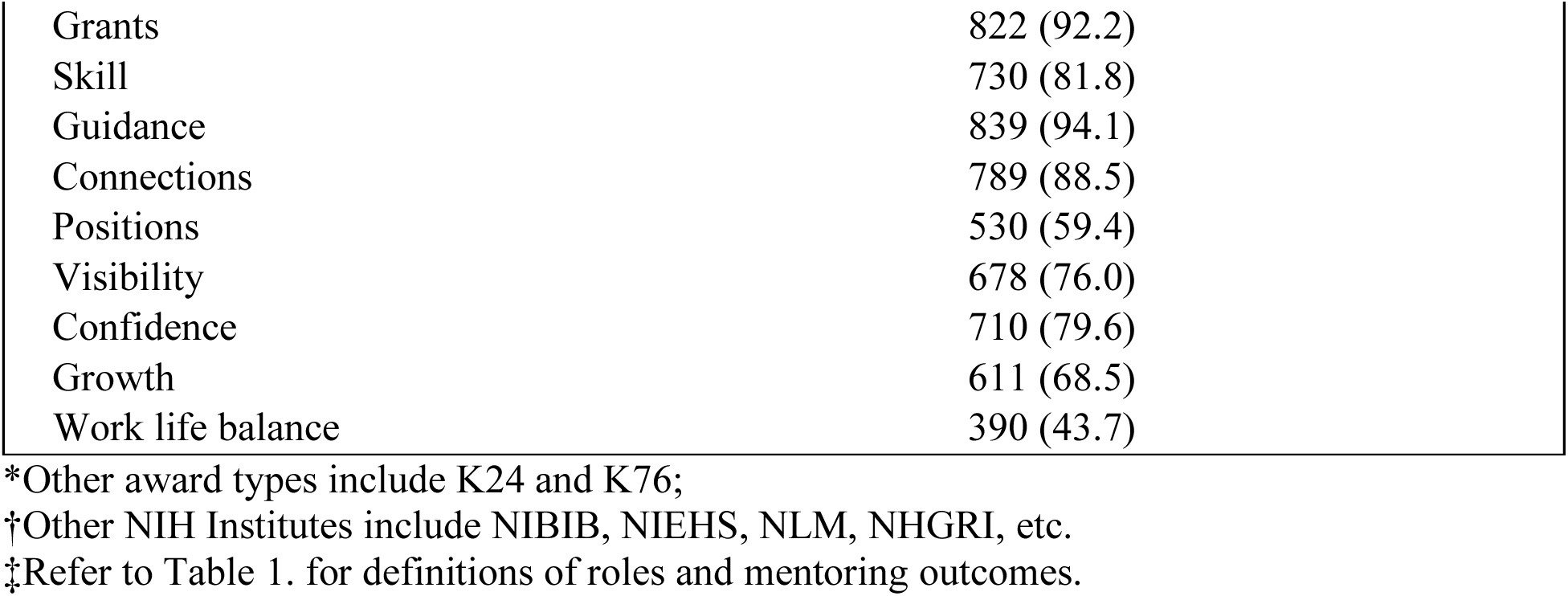
Award and mentor network characteristics of survey respondents who applied for a K-award between 2018-2023 and were awarded (N=892).

### K-awardees have other mentors beyond their primary PI, who play various roles

In terms of mentor characteristics, K-awardees reported having a median of 3 mentors during the time of K-award submission (range = 1-15), including their primary PI (research mentor). After Research mentors, about a quarter of K-awardees have a mentor who primarily played the role of a Coach (24.9%). Slightly less than 20% of K-awardees have mentors whose main roles were of Sponsor and Peer mentors (Table 3). However, when looking at the other (secondary) roles each mentor may be playing, the majority of K-awardees reported having a mentor who play the roles of Connector, Sponsor, and Coach (all >75.0%). For mentoring outcomes (Table 3), most of the K-awardees stated that their mentoring relationships led to Publications and Grants, as well as being able to receive Guidance (all >90.0%). After these categories, the most common mentoring outcomes included new Connections and enhanced Skill, with 88.5% and 81.8% of the K-awardees reported receiving these outcomes from their mentors. The rarest mentoring outcome was receiving strategies to achieve a better Work-life balance (43.7%).

### Mentoring network size is associated with increased variety in mentor roles and mentoring outcomes

Having more mentors was positively associated with having an increased number of distinct mentoring roles (both primary and secondary) in one’s network (coefficient= 0.29, *P*<0.001 for primary, and coefficient = 0.40, *P*<0.001 for secondary). Similarly, having more mentors was also positively associated with receiving higher number of distinct mentoring outcomes (coefficient: 0.40, *P*<0.001). These associations remained true regardless of demographic characteristics.

### Mentor roles are associated with specific types of mentoring outcomes

There were statistically significant differences in the probability of specific mentor roles being attributed to the various mentoring outcomes. The proportions of mentors with a particular role resulting in each mentoring outcome can be seen in Figure 1 (A) and (B). Statistically significant differences in the probabilities calculated by adjusted multi-logistic regression models are presented in Supplemental Table S1. In general, mentors whose primary role is Research have the highest probability of resulting in research-related outcomes including Publications, Grants, and Skills compared to other mentor roles. Coaches had the highest probability of being attributed with Guidance, though not significantly different from those whose main roles are Sponsor, Peer, and Identity mentors. Career-related outcomes, including Connections, Positions, and Visibility, were most likely to be attributed to Connectors and Sponsors compared to any other types of mentors. Finally, psychosocial mentoring outcomes, such as increased Confidence, Growth, and Work-life balance were more likely attributed to mentors whose primary roles were Coaches, Peer, or Identity mentors. However, the differences in the probability of various mentor roles being attributed to specific types of mentoring outcomes were not as stark once all the roles that mentors play, both primary and secondary, were included in consideration (see Figure S1.).

**Figure 1.**
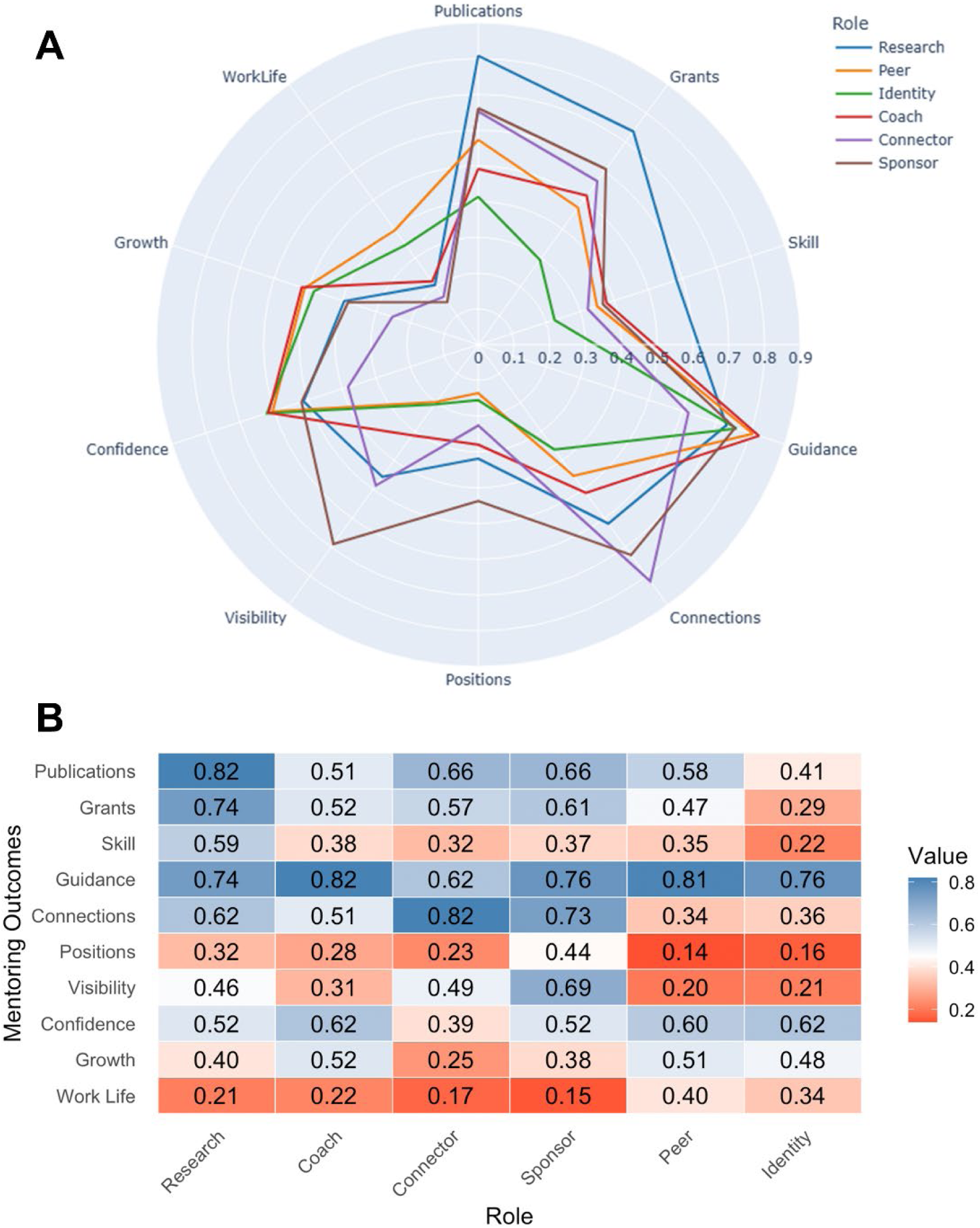
Proportions of mentors with specific primary roles resulting in each mentoring outcome. **(A)** Spider plot displaying the proportion of mentors’ primary roles being attributed to different mentoring outcomes. The numbers by each concentric circle line signify the proportion (e.g., about 80% of mentors whose main role were Research mentors were attributed to the Publications mentoring outcome). **(B)** Heatmap displaying the proportion of mentors’ primary roles being attributed to different mentoring outcomes, with blue signifying higher proportions and red signifying lower proportions.

### Mentoring network characteristics are associated with better K-award scores for those underrepresented in science

While there were no relationships between mentor network size and K-award scores, this relationship is significantly different between those who identify as UR individuals and those who do not (Figure 2), indicating that having more mentors is additively beneficial for UR individuals. Table 4 shows that on average K-award scores are significantly higher (worse) for UR individuals than for their counterparts (mean difference= 4.27, *P=*0.001). However, by having more mentors, UR individuals may be able to catch up. Particularly, according to the model, UR individuals’ scores are reduced by about 1 point for every additional mentor they have.

**Figure 2.**
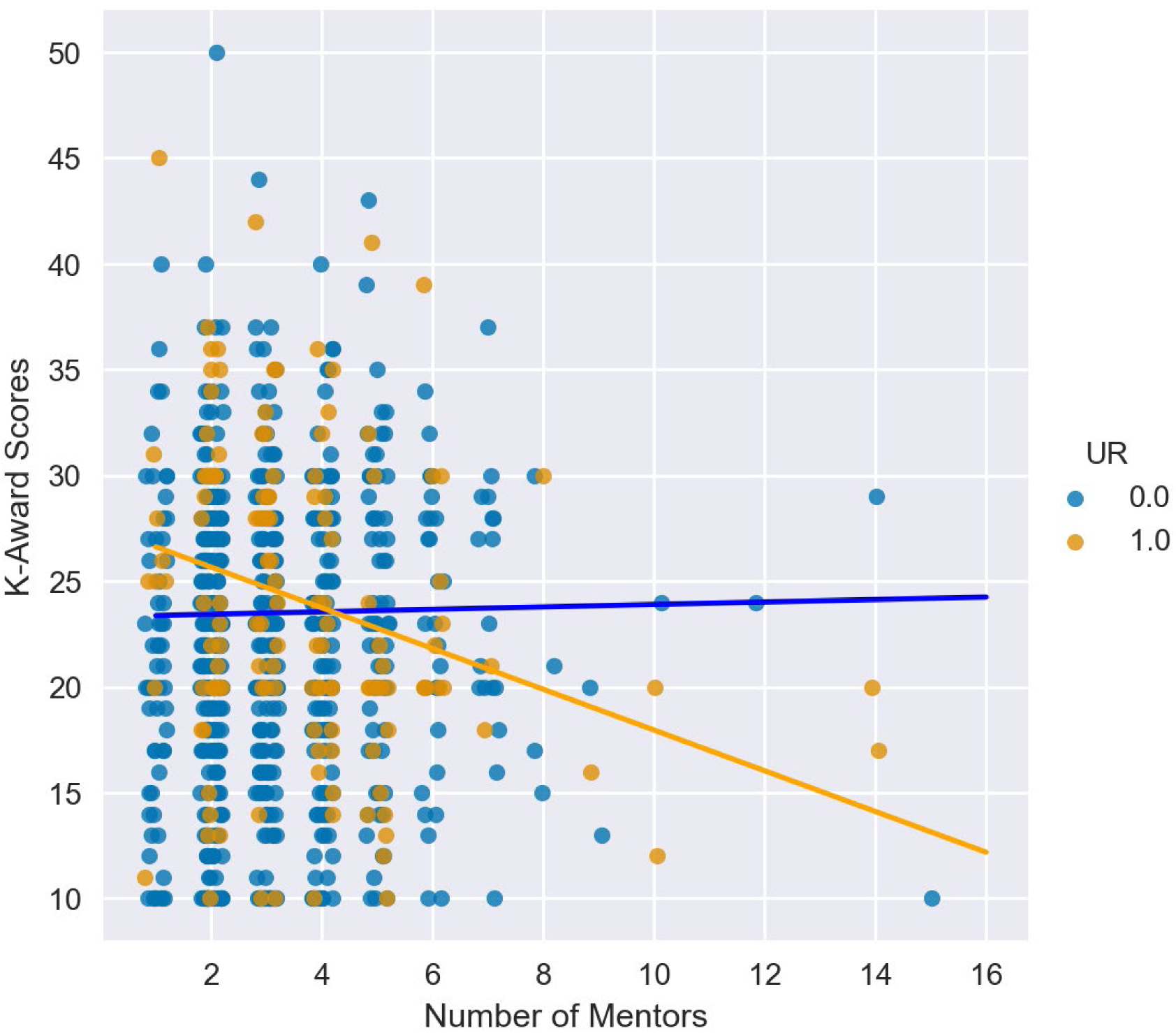
Relationship between number of mentors and K-award scores. Coefficients were obtained from a multilevel linear regression nested by NIH Institute and award type, and further adjusted for gender, first generation status, and density of connections (R2 value= 0.22).

**Table 4.** Coefficients of a mixed linear regression between K-award scores and number of mentors.

| Factor | coefficient | Confidence interval | *P* value |
| --- | --- | --- | --- |
| Number of mentors | 0.06 | -0.33 – 0. 45 | 0.768 |
| Density of connections | -1.69 | -4.66 – 1.28 | 0.264 |
| Sex (male vs female) | -0.26 | -1.17 – 0.65 | 0.571 |
| First generation status | 0.60 | -0.42 – 1.61 | 0.249 |
| UR status | 4.27 | 1.81 – 6.72 | **0.001** |
| UR*number of mentors (interaction) | -1.02 | -1.60 – -0.44 | **0.001** |

To elucidate how having more mentors help UR individuals, additional analyses were conducted. Particularly, we wanted to determine whether there were specific mentor types that were associated with better K-award scores, and how they may be moderated by UR status. We found that the effect of having an identity and peer mentors towards award scores is more beneficial for UR individuals. The models showed significant interaction terms between UR status and having ‘Identity’ (coef= -4.57, *P*=0.023) and ‘Peer’ mentors (coef= -3.44, *P*=0.026).

Similar analysis was done to examine whether UR status moderated the relationship between specific mentoring outcomes and better K-award scores. The models showed that having more ‘Grants’ and ‘Confidence’ mentoring outcomes and K-award scores played a stronger role on award scores for those who are UR individuals (significant interaction terms between UR status and number of ‘Grants’ (coef=-0.96, *P*=0.009) and ‘Confidence’ (coef= -0.95, *P*=0.007) mentoring outcomes).

## Discussion

In this study, we looked at patterns of mentor networks amongst NIH K-awardees and found that 1) typically K-awardees operate within multi-mentor, multi-role networks rather than dyadic mentorship models, 2) mentors can have varying, functionally distinct roles and these roles are associated with specific mentoring outcomes, and 3) mentor network size and composition benefit UR individuals differently than their non-UR counterparts. This is one of the first studies to capture the mentorship networks of postdoctoral trainees across various research fields. We quantified K-awardees’ formal and informal mentors, as well as their mentoring outcomes, allowing us to identify patterns of mentor network structures that may be crucial for scientific advancement. The findings of this study are timely given the increasing concern of attrition of advanced trainees, such as postdocs, from the U.S. (22,23). Understanding how effective mentorship should be structured to support the retention and shape individual scientist’s success is especially critical to ensure the future strength of the scientific workforce (5,45).

Our study suggests that K-awardees, on average, rely on more than just their primary PI, having a median number of 3 mentors. Most of these mentors primarily play the role of Research mentors. This could simply be because most K-awards are considered “Mentored” awards which requires applicants to choose a mentor/mentoring team to apply (46,47). For these awards, there is a strong emphasis on applicants having a strong mentoring team whose collective expertise aligns with and supports the proposed research (46). Having more research mentors may increase the success of K-award applications, since these awards often emphasize the feasibility of the proposed research (48). Exposure to multiple research mentors also impacts trainees broadly; trainees receive wider range of skills, perspectives, and expertise, which, when integrated well into one’s independent research topic, has been shown to be associated with long-term academic success (17,49).

The role of mentorship in K-awardees’ success, however, goes beyond the parting of research expertise. Mentors’ seniority, funding situation, experience in training other K-awardees and/or training grants, and accessibility are all part of the consideration (46,48,50). Mentors are expected to cultivate a plan for their mentees’ career and professional development (47,48). Ripley et al., (2012) sought to understand the importance of having mentors for various training areas (e.g., increased skills in grant writing/manuscript, enhanced research knowledge, networking, etc.) from the perspective of K-awardees and their mentors; they found awardees’ perception of their mentors’ contribution to all areas were lower than that of their mentors, except for networking (50). This indicates the importance of other types of mentoring outcomes, such as increased visibility and/or sponsorship for trainees and how such outcomes may be undervalued by mentors. In fact, our study found that as K-awardees expand their network, their mentors tend to play different roles, and they receive a wider variety of mentoring outcomes. These findings suggest increasing mentorship network size brings about the various mentoring outcomes needed to maximize research success and professional development for postdoctoral scientists.

As one expands the size of their network, one should proactively seek mentors that are most conducive to result in desired mentoring outcomes. Our study found for example, research-related mentoring outcomes, such as Publications, Grants, and research Skills were attributed to Research mentors significantly more than to any other mentor roles. Another study found similarly that K-awardees’ primary research mentors more commonly resulted in research-related outcomes compared to other types of outcomes (e.g., job promotion, clinical experience, etc.) (50). It is quite common for postdocs to report dissatisfaction with the amount of support and guidance they receive from their research mentors on career-related issues (15,40). Instead, our study suggests that career-related mentoring outcomes such as new Connections, Positions, and increased Visibility may be best directed to mentors who are Connectors and Sponsors. While they differ in function, both have been reported in the literature to be instrumental for career advancements. Sponsors are typically more senior researchers/clinicians who actively and visibly use their credentials and reputation to advocate for and advance trainees’ careers, in spaces where opportunities may have otherwise been absent (51–53). Connectors are those who leverages their network to ‘connect’ mentees with relevant individuals (31). These connections often work powerfully for postdocs who are trying to increase their visibility and/or secure new positions (54,55). Generally in both academic and medicine settings, an overreliance on dyadic (one-on-one) mentorship relationships still persist (6,56). One mentor, however, is unlikely to meet the various needs a mentee has (57). This study affirms that one can much more efficiently receive holistic mentoring, if one is strategic about the types of mentors they seek.

Among the list of mentoring outcomes, psychosocial related needs, such as Growth and Work-life balance, were the least met among K-awardees. However, these types of mentoring needs should not be ignored. Rising rates of burnout and a lack of work-life balance continue to impact trainees’ satisfaction and desire to advance in the academic and clinical pipelines (16,18,20,21,58). In this study, we found that increased Confidence was typically obtained from Coaches, Peer, and Identity mentors. In the context of skill improvement such as in grant-writing, having coaches can significantly improve one’s confidence (59). Improved Work-life balance and personal Growth were mostly attributed to Peer and Identity mentors. The importance of these types of mentors have been widely discussed elsewhere; informal and formal peer mentors can effectively create a sense of belonging, increase confidence, and improve scientific identity among mentees (34,38,60). Likewise, Identity mentors (those who have shared identities, for example, being a first-generation trainee and/or those who excel in culturally-responsive mentoring practices) can provide insights into the unique challenges that individuals may face based on their identities (15,28,39). Consequently, identity mentors are also often attributed to improving confidence, affirming belonging, and increased mentoring relationship satisfaction (38,61). The psychosocial needs Peer and Identity mentors often fulfill should not be undervalued as they carry long-term implications for career retention in STEMM (5,42).

Our study found that having a larger mentor network additively benefits UR compared to their non-UR counterparts. This could point to the fact that UR individuals generally find it more difficult to have all of their mentoring needs met from their primary mentors, with many feeling that they have to find external mentors to support their success (15,40). Prior research has documented that UR individuals encounter unique challenges in academic environments, including limited social capital, fewer identity-similar role models, and increased experiences of bias, discrimination, and social isolation (4,5,15,38,56). Identity mentors, those who share similar characteristics and/or those trained to recognize and address the unique experiences of UR mentees, can play a critical role in supporting their success (38,62). They, along with Peer mentors have been found to boost confidence and effectively cultivate scientific identity for UR individuals; which is particularly important given that UR individuals are less likely to see others like them in faculty and/or leadership positions (15,41,60,63). This is corroborated by our findings where having Peer and Identity mentors, and receiving more confidence significantly benefited UR individuals’ K-Award scores more than their non-UR counterparts. In addition to above, Peer mentors may remove the difficulty of navigating power dynamics and can provide UR individuals advice and sample grant materials that help boost the likelihood of K-Award applications (55,59). The presence of Peer and Identity mentors in one’s network may benefit UR individuals in complementary ways and overall, reduce disparities in grant success (64).

This study is not without limitations. The sample was obtained through convenience sampling, which could lead to non-response bias, where those who chose to not respond to the survey is significantly different than those who did. Our sample have more women (62.9%) compared to the population (range of 52-58% between 2018-2023) based on data released by the NIH Division of Statistical Analysis and Reporting (65) but this could be because women tend to be more willing to complete surveys compared to men (66). While we had initially recruited individuals who applied for the K-Award, the majority of our respondents were those who received the K-Award. Those who had positive mentoring relationships and were successful in their applications may be more motivated to complete the survey. Due to the imbalanced sample, we could not compare whether the network characteristics of those who received the K-award differ from those who did not. Though we recommend future research to pursue such inquiry. Despite these limitations, our recruitment strategy still allows us to obtain a sample that is relatively representative of the national population of researchers who were awarded a research grant by NIH (67). Our study is also one of the first to collect data on both formal and informal mentors going beyond co-authors and primary PIs as others typically have done (17,49). Hence, our data reflects a truer reality of the relationships that surround current postdocs, especially as virtual and cross-institutional mentoring relationships are increasingly common now (68,69).

Based on our findings, we strongly urge mentoring networks to be incorporated as part of scientific training for advanced STEMM trainees, with a focus on growing one’s network size and diversifying the roles that one’s network play. As networks expand, trainees may require tools and institutional support to effectively manage multiple mentoring relationships, coordinate expectations, and sustain mentee agency. The development of structured mentor-mapping frameworks, digital platforms, and evidence-based guidance for maintaining functional mentoring networks may therefore represent an important next frontier in mentorship science. With growing concerns for scientific talent attrition in the U.S., we encourage further research to increase understanding on the different mentoring structures that may be most effective for individuals of various identities and fields, including international trainees. Future research would move beyond cross-sectional analysis and explore a longitudinal approach to understand how mentorship networks evolve over time and how changes in network size, role diversity, and composition relate to subsequent career milestones. In addition, intervention-based studies are needed to test whether intentional diversification of mentorship networks improves trainee outcomes. Programs such as Structured Professional Networks for Successful Outcomes in Research (70) provide a promising model to experimentally evaluate whether scaffolding network development through mentor (role) mapping, sponsor identification, and identity-affirming mentorship can strengthen career progression. Additionally, we acknowledge there are other mitigating factors that may influence whether trainees will have their goals met by their mentors such as relationship strength and meeting frequency; we suggest such variables to be collected and evaluated in future mentoring interventions.

## Acknowledgments

### Funding

National Institute of General Medical Sciences 5R35GM151017-03 (WML)

### Author contributions

Conceptualization: WML

Methodology: FJS, WML

Investigation: FJS, WML

Visualization: FJS, EH

Supervision: WML

Writing—original draft: FJS, EH

Writing—review & editing: FJS, EH, WML

### Data and materials availability

Raw data is deposited and available at https://www.openicpsr.org/openicpsr/project/251270/version/V1/view. Identifiable variables were removed and/or de-identified to protect the anonymity of the survey respondents.

Explanations on how data was prepared prior to submission to the ICPSR repository is available in the Documentation file, also available at the same link.

