## Supplementary Table S1 for "How Mentoring Networks Shape Early-Career Grant Success: Evidence from NIH K-awardees"

**Table S1. Average proportion of mentors with a specific primary role being attributed to each type of mentoring outcomes.** Different letters signify that the proportions are significantly different from one another.

| Mentoring Outcomes | Mentor roles | | | | | |
|---|---|---|---|---|---|---|
| | Research | Coach | Connector | Sponsor | Peer | Identity |
| Publications | 0.82[a] | 0.51[c] | 0.66[b] | 0.66[b] | 0.58[bc] | 0.41[c] |
| Grants | 0.74[a] | 0.52[b] | 0.57[b] | 0.61[b] | 0.47[bc] | 0.29[c] |
| Skill | 0.59[a] | 0.38[b] | 0.32[b] | 0.37[b] | 0.35[b] | 0.22[b] |
| Guidance | 0.74[bc] | 0.82[a] | 0.62[c] | 0.76[abc] | 0.81[ab] | 0.76[abc] |
| Connections | 0.62[b] | 0.51[c] | 0.82[a] | 0.73[ab] | 0.34[c] | 0.36[c] |
| Positions | 0.32[b] | 0.28[b] | 0.23[bc] | 0.44[a] | 0.14[c] | 0.16[bc] |
| Visibility | 0.46[b] | 0.31[c] | 0.49[b] | 0.69[a] | 0.20[c] | 0.21[c] |
| Confidence | 0.52[bc] | 0.62[a] | 0.39[c] | 0.52[abc] | 0.60[ab] | 0.62[abc] |
| Growth | 0.40[b] | 0.52[a] | 0.25[c] | 0.38[abc] | 0.51[a] | 0.48[ab] |
| Work life | 0.21[bc] | 0.22[bc] | 0.17[bc] | 0.15[c] | 0.40[a] | 0.34[ab] |
