## Supplementary Figure S1 for "How Mentoring Networks Shape Early-Career Grant Success: Evidence from NIH K-awardees"

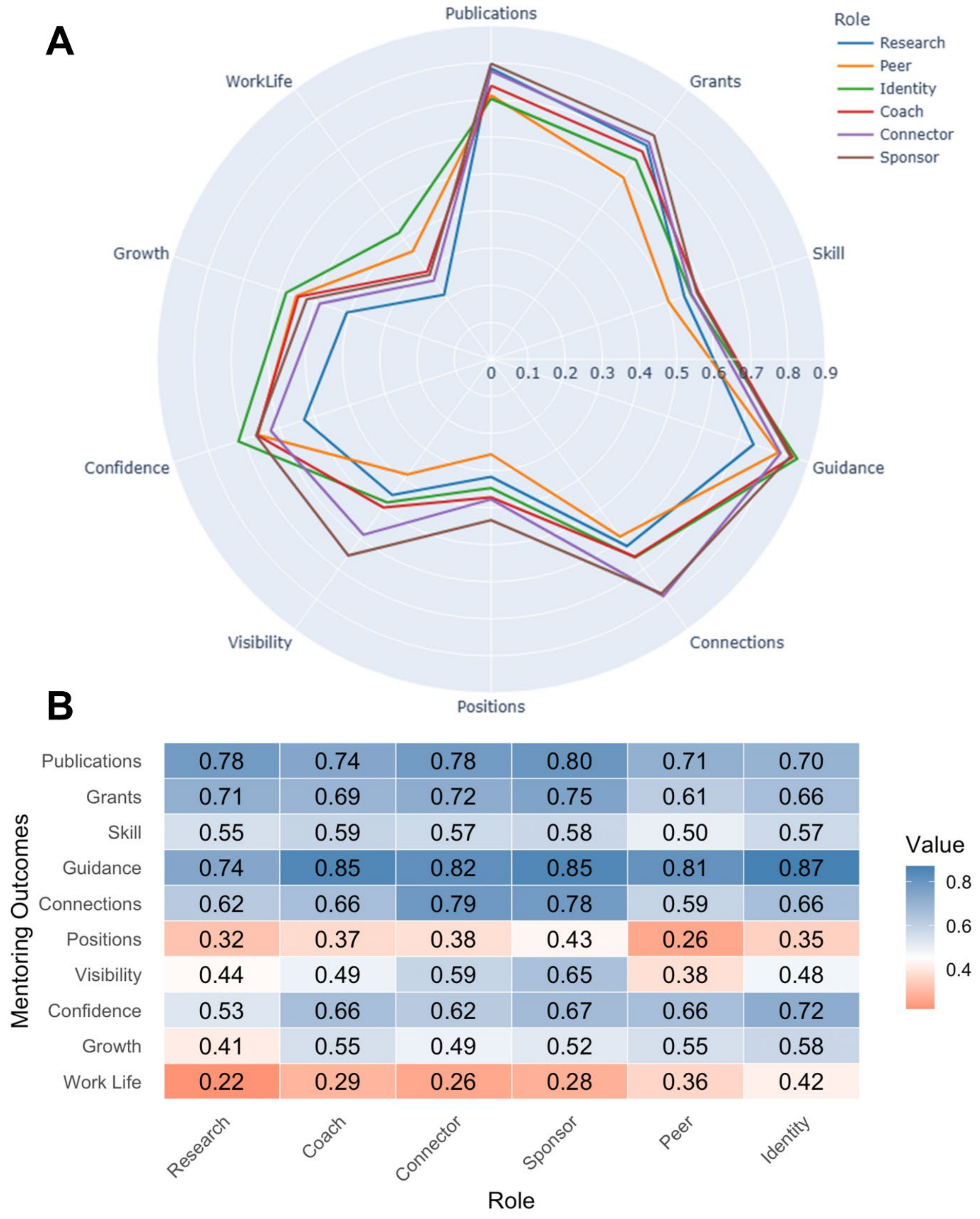


**Fig. S1. Proportions of mentors with specific roles (both primary and secondary) resulting in each mentoring outcome.** (A) Spider plot displaying the proportion of mentors' roles being attributed to different mentoring outcomes. The numbers by each concentric circle line signify the proportion (e.g., about 80% of mentors who had a "Research mentors" role were attributed to the “Publications” mentoring outcome). (B) Heatmap displaying the proportion of mentors’ roles being attributed to different mentoring outcomes, with blue signifying higher proportions and red signifying lower proportions.
